# Hyperoxia Inhibits the Growth of Mouse Forebrain Oligodendrocyte Progenitors

**DOI:** 10.1101/2021.01.20.427261

**Authors:** Lisamarie Moore, Lauren E. McLane, Stacey Wahl, Isis M. Ornelas, Teresa L. Wood, Peter Canoll, Steven W. Levison

## Abstract

NG2 chondroitin sulfate proteoglycan positive oligodendrocyte progenitor cells (OPCs) reside throughout the brain. They divide asymmetrically and differentiate into myelinating oligodendrocytes throughout adulthood. OPCs have been successfully isolated from rodents using several techniques including magnetic beads, immunopanning and exploiting differential centripetal adhesion. Whereas rat OPCs are relatively simple to propagate *in vitro*, it has been difficult to expand mouse OPCs. Therefore, we evaluated the effects of oxygen levels, growth factors and extracellular matrix components to produce a simple and reproducible method to prepare large numbers of nearly homogenous cultures of primary mouse OPCs from postnatal day 0-2 mouse telencephala. Using the McCarthy and de Vellis mechanical separation method OPCs were separated from mixed culture of glial cells. When the OPCs were plated onto fibronectin coated tissue culture plates in a biochemically defined medium that contained fibroblast growth factor-2 (FGF-2) and platelet derived growth factor AA (PDGFAA), and they were maintained in a standard tissue culture incubator, they proliferated very slowly. By contrast, mouse OPCs doubled approximately every 7 days when maintained in a 2% oxygen, nitrogen buffered environment. After 3 passages, greater than 99% of these OPCs were NG2+/PDGFRα+. In medium containing only FGF-2, mouse OPCs progressed to late stage OPCs whereupon A2B5 expression decreased and O4 expression increased. When these cells were differentiated between passages 1 and 3, the majority of the OPCs differentiated into MBP+ mature oligodendrocytes However, cells that were repeatedly passaged beyond 4 passages progressed to a late O4+ OPC (even with mitogens present) and when differentiated by mitogen removal a minority of the OPCs differentiated into MBP+ cells. These studies reveal significant differences between mouse and rat OPCs and an inhibitory role for oxygen in mouse OPC proliferation.

## Introduction

One of the last cells to develop from the neuroepithelium of the central nervous system (CNS) is the oligodendrocyte, which become the myelinating glia of the CNS, and play a critical role in facilitating saltatory conduction of neuronal action potentials as well as supporting axonal survival [1,2]. Oligodendrocytes are concentrated in the white matter but are also found extensively throughout the brain parenchyma. Most adult OPCs will remain immature proliferative cells while a subpopulation can differentiate into mature oligodendrocytes to replace dying or dead cells throughout the lifespan [3,4].

Oligodendrocyte progenitor cells (OPCs) proliferate and migrate throughout the CNS beginning in late embryonic development, They pass through several distinct stages as they mature that can be distinguished using antigens that are expressed at distinct stages [5,6]. Platelet-derived growth factor receptor alpha (PDGFRα), a tyrosine kinase receptor, is expressed by early stage OPCs. OPCs that express PDGFRα generally have a bipolar morphology. These cells also express tetrasialogangliosides that are recognized by the A2B5 monoclonal antibody, which are also found on multiple CNS cell types, but when expressed by an OPC identifies it as an early progenitor cell [7-9]. Another useful antigen for the early OPC is NG2, a chondroitin sulfate proteoglycan [3]. In addition, NG2 and A2B5 expressing cells maintain the capacity to terminally differentiate into astrocytes as well as oligodendrocytes [3]. These markers are useful in identifying immature OPCs but must be used with care as both can label other neural precursor cell populations.

As OPCs mature into oligodendrocytes they start to express antigens recognized by O4, a monoclonal antibody that binds cell surface pro-oligodendroblast antigen (POA) and sulfatide, which are found on late-stage OPCs and post-mitotic oligodendrocytes. It serves as a more reliable marker for OPCs committed to an oligodendroglial fate than A2B5 or NG2 alone [3,4,10]. O4 expressing cells typically have a multipolar morphology. Finally, mature myelinating oligodendrocytes express myelin specific antigens such as galactocerebroside (GalC) and myelin basic protein (MBP). O1 is a monoclonal antibody that binds to galactocerebroside and identifies pre-myelinating oligodendrocytes that no longer maintain their proliferative potential but are transitioning to a terminally differentiated oligodendrocyte [11,12]. MBP is one of the primary protein constituents of myelin and a marker for mature oligodendrocytes [4].

The isolation and *in vitro* study of OPCs from both the developing and adult CNS has provided critical insights into our understanding of injury and potential repair after white matter damage from demyelinating diseases like multiple sclerosis, or traumatic injury to the brain and spinal cord or brain tumors. Numerous oligodendroglial cell lines have been developed, such as the CG4 cell line [13], but the molecular changes that have occurred with transformation render these cells inadequate for many studies of oligodendrocyte function and myelination.

Several methods for isolating rat OPCs from the CNS have been described, such as immunopanning [6,14-16], fluorescence-activated cell sorting (FACS) and magnetic cell sorting (MACS) by exploiting cell surface-specific antigens [6,16-18]. However, most of these methods use A2B5 or NG2, and thus select for cells that are and are not committed to the oligodendroglial lineage. Other methods have included differential gradient centrifugation [19-21], formation of oligospheres [22] and lastly the shaking method based on differential and mitotic adherent properties of glia [23-25]. This method permits the separation of rat OPCs from an astroglial bedlayer by centripetal forces. This method has been very successful for isolating rat OPCs, but mouse OPCs do not proliferate in the same manner as rat OPCs after isolation. As most transgenic and knockout studies are carried out in mice, there is a crucial need to develop a simple procedure to produce ample amounts of OPCs at high purity from mice.

In this study we describe a reliable and reproducible method for the isolation, purification and expansion of a highly enriched preparation of OPCs in sufficient numbers for experimentation from mice. The OPCs may be passaged up to three times to obtain early OPCs with minimal phenotypic drift while increasing the purity of the cultures. After approximately 6 repeated passages, these OPCs resemble the phenotype of adult OPCs. These mouse OPCs maintained the ability to differentiate with limited astrocyte differentiation. Overall this method will facilitate future studies of OPC differentiation and oligodendrocyte function.

## Materials and Methods

### Mixed Glial Cell Cultures

Cultures were established from C57Blk/6 mouse telencephala, from postnatal day 0 (P0) to day 2 (P2). Mice were decapitated under sterile conditions and their brains were placed into chilled phosphate buffered saline (PBS) with 0.6% d-glucose and 2 mM MgCl_2_ (PGM) on ice blocks. The meninges were carefully removed using fine forceps and the block of tissue excised between the olfactory bulbs and the cerebellum. This block was transferred to fresh PGM and the isolated tissue was gently minced with forceps. The tissue was then transferred to conical tubes using a large bore P1000 tip or a 10mL pipette and centrifuged at 200 x g for 10 minutes. The pellet was enzymatically dissociated using a pH-balanced solution containing 1 ml trypsin (2.5% stock) and DNase I (0.02mg/mL) dissolved into 9 mL of minimum essential media (MEM). The tissue was digested for 5-7 minutes @ 37° C with manual agitation several times during incubation. An equal volume of medium supplemented to 10% fetal bovine serum (FBS) was added to quench the trypsin. The mixture was triturated for several cycles using a 10mL pipette, adding additional media during later cycles. The cell suspension was passed through a 100 μm cell strainer to eliminate clumped cells from the final mixture. Then the cells were centrifuged at 400 x g for 8 minutes. Finally, the supernatant was removed and passed through a 40 μm cell strainer. Viable cells were counted and cells were plated at 2×10^5^ cells/cm^2^ in MEM-10C media [High glucose MEM medium containing 10% (v/v) pre-screened^1^ FBS (Atlantic Biologicals), 200mM L-Alanyl-L-Glutamine (CellGro), 1% Penicillin-Streptomycin solution (Corning-CellGro) and 0.6% d-glucose (Sigma-Aldrich). Cells were cultured in twist-cap Falcon tissue culture straight neck T75 flasks (Becton Dickinson #3024) for 10 −12 days with the first medium change 4 days after the initial plating and then every other day onwards. All animal work was performed according to an approved Rutgers NJMS Institutional Animal Care and Use Committee (IACUC) protocol #10012.

### Isolation and Purification of Mouse Oligodendrocyte Progenitors

Once the bed layer of the mixed glial cultures was confluent the mouse OPCs were isolated using the well-established mechanical separation technique [23] with modifications to improve cell survival as described previously [26]. The twist-caps on the T75 Falcon flasks were tightened, and then the flasks were subjected to centripetal shaking @ 260 rpm for 1.5 hours @ 37°C to remove lightly adhered microglia. Flasks were gently agitated manually to remove any remaining adhered microglia, followed by supernatant removal. Cells were rinsed twice with supplemented MEM and then placed back in the incubator in MEM-10C for at least 1 hour to equilibrate with 5% CO_2_ @ 37°C. The twist-caps were tightened once more and the cells shaken overnight for 18hrs @ 260 rpm to remove the attached OPCs while leaving the more adherent astrocytes behind. The flasks were gently agitated manually to further dislodge any remaining OPCs. The supernatant was removed and very slowly passed through a 20 μm nitex mesh (Amazon.com) affixed to a sterile beaker to remove any clumps of astrocytes that may have been dislodged. Unsupplemented MEM was added to the flask twice to gather any remaining OPCs. The cells were then transferred to a conical tube and centrifuged at 400 x g for 5 min. Then cells were resuspended in N2B2 media [DMEM:F12-HEPES supplemented to 1X (v/v) B27 supplement (Gibco), 1% (v/v) Penicillin-Streptomycin solution (Corning-CellGro) and 0.5% (v/v) prescreened FBS (Atlanta Biologicals). The cells were plated onto a 100 mm bacteriological plastic dish for 30 minutes @ 37°C to remove contaminating microglia [16]. Bacteriological plastic dishes were rinsed twice with N2B2 to collect the nonadherent OPCs while leaving behind any attached microglia. The supernatant was centrifuged at 400 x g for 5 minutes. Viable cells were counted and resuspended in N2S [N2B2 medium supplemented with 30% B104 neuroblastoma conditioned medium (B104 CM) [26], or 10 ng/mL recombinant rat platelet-derived growth factor (PDGF-aa) (R&D Systems) and 10 ng/mL FGF-2 (Peprotech) with 1 ng/mL heparin sulfate (Sigma-Aldrich) (proliferation media). OPCs were plated onto 100 mm tissue culture plates that had been previously coated with 0.5 μg/cm^2^ fibronectin (MW > 440k, BD Biosciences) for 2 hours @ 37°C. The OPCs were seeded at 3×10^4^ cells/cm^2^ and maintained in either 2% O_2_, 5% CO_2_, 93% N_2_ or 20% O_2_, 5% CO_2_, room air @ 37° C. Half of the medium was changed every other day until the cells were approximately 80% confluent.

### Amplification and Differentiation of Mouse Oligodendrocyte progenitors

OPCs were gently removed from the fibronectin-coated plates using solution containing 44 U/mL papain (Worthington) and DNase I in MEM with 20 mM Hepes, 5 mM EDTA and 14 mM L-cysteine. The solution was swirled over the cells and then aspirated to leave a thin film of solution. The cells were returned to the incubator for 7 min. Plates were agitated by hitting vigorously. Once the cells were detached, 5 mL of N2B2 was added and the cells were collected by centrifugation at 400 x g for 8 min. The pellet was resuspended and viable cells were plated in N2S on fibronectin-coated flasks at no less than 2.5×10^4^ cells/cm^2^. Again, half media changes were performed every other day until the cells were 80% confluent and ready to be passaged. To differentiate the OPCs, the mitogens were removed from the media and the cells were placed into N2B2 supplemented with 1.95 μg/mL triiodothyronine (T3) (Sigma) [27,28], 1 ng/mL recombinant rat CNTF (R&D Systems) [29], and 1 ng/mL TGFß1 (R&D Systems) [30] for 5 – 7 days_[LMM1]_. Then stained for mouse anti-O4 (1:4, supernatant), mouse anti-O1 (1:4, supernatant), rat anti-PLP (1:1000, gift from Wendy Macklin), and mouse anti-MBP (1:250, Covance), rabbit anti-Olig2 (1:100, Milipore), and rat anti-GF2.2 (1:4, supernatant).

### Immunofluorescence

Cells were replated onto fibronectin-coated chamber slides (LabTek) and maintained for 24 to 72 hours in N2S (proliferation media) or supplemented N2B2 (differentiation media). To identify astrocytes, oligodendrocytes and microglia, cells were stained live for mouse anti-O4 (1:5, supernatant), mouse anti-A2B5 (1:4, supernatant), rabbit anti-NG2 Chondroitin Sulfate Proteoglycan (1:50 Millipore) and rat anti-PDGFRα (1:100 BD Pharmigen) for 1 hour at room temperature, rinsed and fixed with 4% paraformaldehyde (PFA) and sometimes followed by 100% ice chilled methanol. The cells were incubated with appropriate fluorochome-conjugated secondary antibodies for 1 hour at room temperature. The cells were then blocked, permeabilized and incubated with mouse or rabbit anti-Olig2 (1:100, Millipore), rat anti-CD11b (1:50, BD Pharmingen), GF2.2 (1:4, supernatant), rabbit anti-Sox (1:500), and rabbit anti-Ki67 (1:750, Vector). The cells were washed and further incubated with secondary antibodies. Some cells were counterstained with 1 μg/ml DAPI (Sigma) for 5 minutes. For differentiation studies, cells were stained for markers of the oligodendrocyte lineage with markers MBP (1:250, Covance), PLP (1:3, supernatant), and O1 (1:3, supernatant) followed by appropriate secondaries. All secondary antibody combinations were carefully examined to ensure that there was no bleed through between fluorescent dyes or cross-reactivity between secondary antibodies. No signal above background was obtained when the primary antibodies were replaced with pre-immune sera. All secondaries were purchased from Jackson ImmunoResearch.

Stained cells were washed thoroughly and mounted with Fluorogel (Electron Microscopy Sciences) and allowed to dry overnight. Immunoreactive cells were visualized using an Olympus Provis AX70 microscope and images of stained cells were collected using a Q-imaging CCD camera interfaced with IP Lab scientific imaging software (Scanalytics). Labeled cells in at least 4 random (nonadjacent) fields per well were counted under 20X or 40X objective and a total of 4 wells per independent group were evaluated with at least 100 cells counted based on DAPI staining to evaluate the purity of the isolation and amplification technique described above.

### Surface Marker Analysis by Flow Cytometry

Cell surface expression was assayed by flow cytometry according to the following protocol. OPCs were removed from the tissue culture plates gently with 0.2 Wünsch unit (WU)/ml of Liberase DH (Roche) and DNase I in PGM at 37°C for 5 min. An equal volume of PGM plus DNase I without Liberase was added and the plates were vigorously agitated. Detached cells were collected by centrifugation at 400 x g for 5 min. Cells were resupended in 10 mL of PGB [PGM supplemented with 2 mg/mL fraction V of Bovine Serum Albumin (Fisher)] and centrifuged for 5 min at 400 x g. Cells were dissociated by repeated trituration and then viable cells were counted by hemocytometer. Cells were segregated into 1×10^6^ cells per experimental group (stained, isotype control, live/dead and unstained). Prior to staining, cells were resuspended in 50 μL of FcR block in PGB (1:50, BD Pharmigen). All staining was performed in 0.65 μL eppendorf tubes using a total volume of 150 μL. For surface marker analysis, cells were incubated with unconjugated mouse O4 for 25 min on ice. Cells were washed and then incubated with anti-mouse IgM-PerCP-eFluor 710 (1:200; eBioscience) in 1:4 rat serum for 25 minutes on ice. Cells were then blocked with nonspecific IgMs to bind all available epitopes on the secondary, washed again, and then incubated with unconjugated rabbit anti-NG2 and mouse anti-Glast (1:11; ACSA-1, Milltenyi). After washing with PGB, cells were incubated with appropriate secondaries [goat anti-rabbit IgG Alexa Fluor 700 (1:100; Molecular Probes) and Strep-APC-eFluor 780 (1:160; eBioscience)] along with directly conjugated antibodies CD133-APC (1:50; 13A4, eBioscience), Lewis-X-V450 (1:20; MMA, BD Bioscience), CD140a-Pe (1:400; APA5, Biolegend) and 1:3200 DAPI solution with 1:4 goat serum for 25 min. To restrict our analysis to living cells, we used a live/dead cell exclusion technique that identified DAPI positive cells as dead and DAPI negative cells as alive. Cells were washed with PGB by centrifugation at 500 x g and sometimes lightly fixed with 1% ultrapure formaldehyde in PBS w/o Mg^2+^ and Ca^2+^. Whenever possible, the cells were analyzed immediately after completing the staining protocol. All sample data were collected on the BD LSR II (BD Biosciences Immunocytometry Systems) with excitation lasers at 350, 407, 488, and 633 nm and bandpass filters for APC (660/20), V450 (440/40), Pe (575/26), Alexa 488 (530/30), Alexa700 (710/20), APC-eFluor 780 (780/60), PerCp-eFluor 710 (695/40) and DAPI (440/40). Gates were set based on matching isotype controls from the same manufacturer as the purchased primary antibodies. Afterwards, the acquired data were analyzed by FlowJo software (Tree Star).

### Statistical Analyses

Results were analyzed for statistical significance using a parametric student’s *t* test. Error bars represent SEMs. Comparisons were interpreted when significance was associated with *p* < 0.05.

## Results

### OPCs are mechanically separated and purified from a mixture of glial cells prepared from neonatal forebrain cultures

Mixed glial cell cultures were prepared from cells dissociated from the telencephala of P0-P2 C57BL/6 mice and plated onto T75 flasks in Eagle’s MEM culture medium supplemented to 0.6% glucose and 10% FBS. These cultures were maintained *in vitro* in standard tissue culture incubators until they reached confluency, which typically required 10-12 days (Figure 1A). At confluency the cultures were comprised of phase bright microglia (Figure 1A, blue arrows), phase dark OPCs (Figure 1A, orange arrows) on top of a bed layer of flat adherent astrocytes and fibroblasts. Cultures enriched in OPCs were produced by removing these precursors using centripetal forces as originally described by McCarthy and de Vellis [23]. The mixed cultures were placed onto a rotary shaker for an hour at 260 rpm to remove the loosely attached microglia (Figure 1A1), which was followed by an 18 hour shake to remove the majority of the OPCs (Figure 1A2).

**Figure 1.**
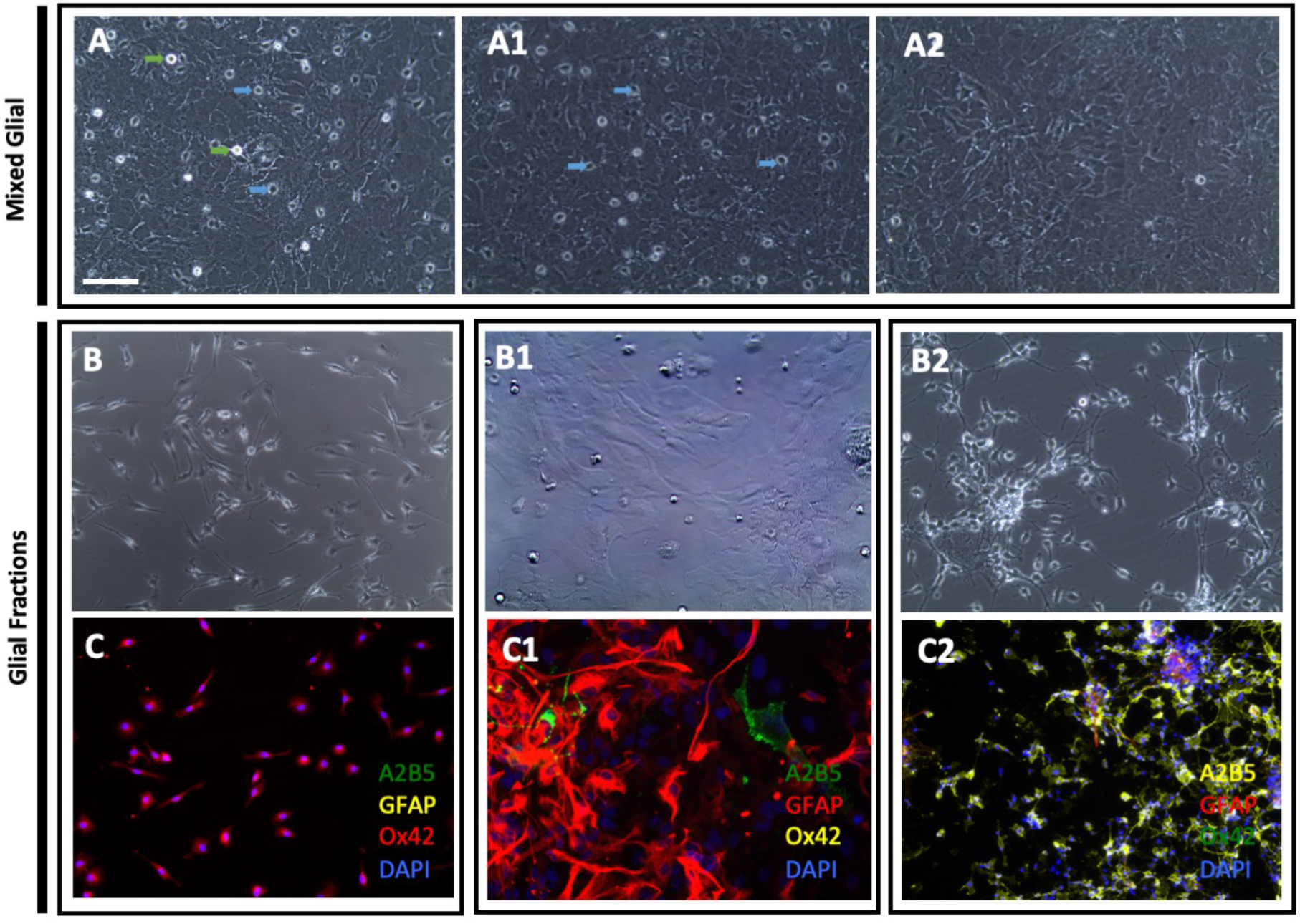
Separation of OPCs from neonatal mouse forebrain mixed glial cell cultures. Mixed glial cell cultures were prepared from dissociated P0 mouse telencephala cultured in MEM-10C (A). Phase bright cells were predominantly microglia (green arrows) and phase dark cells were predominantly OPCs (blue arrows). Flasks were shaken at 260 rpm for 1 h to remove microglia and other loosely attached cells and then shaken a second time at 260 rpm for 18 hours to remove OPCs. Representative phase contrast images of the cultures after the first mechanical shake (A1) and the second shake (A2). Representative phase contrast images of the cells obtained from the first mechanical shake (B), the adherent bed layer after the second shake (B1) and the cells obtained from the second shake (B2). Fractions were stained for A2B5 GFAP, OX42 and DAPI (C1-C2). Scale bar represents 100μm.

The fractions produced by this differential adhesion method were stained for markers of the microglial (Ox42), oligodendrocytic (A2B5) and astrocytic (GFAP) lineages (Figure 1C-C2). The microglia enriched fraction was comprised of predominately elongated amoeboid shaped cells that stained positive for Ox42. There were a few OPCs and astrocytes detected in this fraction (data not shown). The bed layer was comprised mainly of GFAP+ astrocytes with another smaller population of cells that were later identified as αSMA+ fibroblasts (data not shown). This fraction also contained sparse microglia and OPCs (data not shown). The OPC containing fraction was passaged as described below, to produce a more homogeneous cell population as passaging removed more firmly adherent cells, like microglia and astrocytes, while enriching for the OPCs. The purity of the enriched OPC fraction after three serial passages achieved 96.48%±3.6% positive bipolar immature OPCs with a small percentage of contaminating astrocytes (3.60%±3.4) and microglia (0.02%±0.003).

### Large quantities of OPCs are obtained in vitro when cells are cultured in media supplemented with B104 and FGF-2 in physiological oxygen (2% O_2_)

Initially, the enriched OPC fraction was grown in a medium supplemented with 10 ng/mL PDGFaa + 10 ng/mL FGF-2 and the cells were plated onto poly-d-lysine (PDL) coated plates as this method has been used effectively to amplify rat OPCs [31-34]. Under these growth conditions, the OPCs developed elongated processes and were so firmly affixed to the culture plates that they could not be efficiently passaged (Figure 2A). Therefore, we modified the culture medium, supplementing it with B104 neuroblastoma condition media (B104CM) as this medium had been previously reported to support rat OPC growth similar to PDGF and FGF-2 [26]. B104CM has been reported to contain a high molecular weight, heparin binding polypeptide growth factor that stimulates the proliferation and survival of rat OPCs (Bottenstein patent) [35-37]. Although papain effectively detaches viable rat OPCs from a PDL coated surface, it was ineffective for detaching the mouse OPCs; therefore, the culture substratum was coated with fibronectin. With these modifications to the protocol, the cells appeared healthier and were easily detached using papain. However, the cells grew very slowly, generating insufficient cell numbers for future experimentation (Figure 2B).

**Figure 2.**
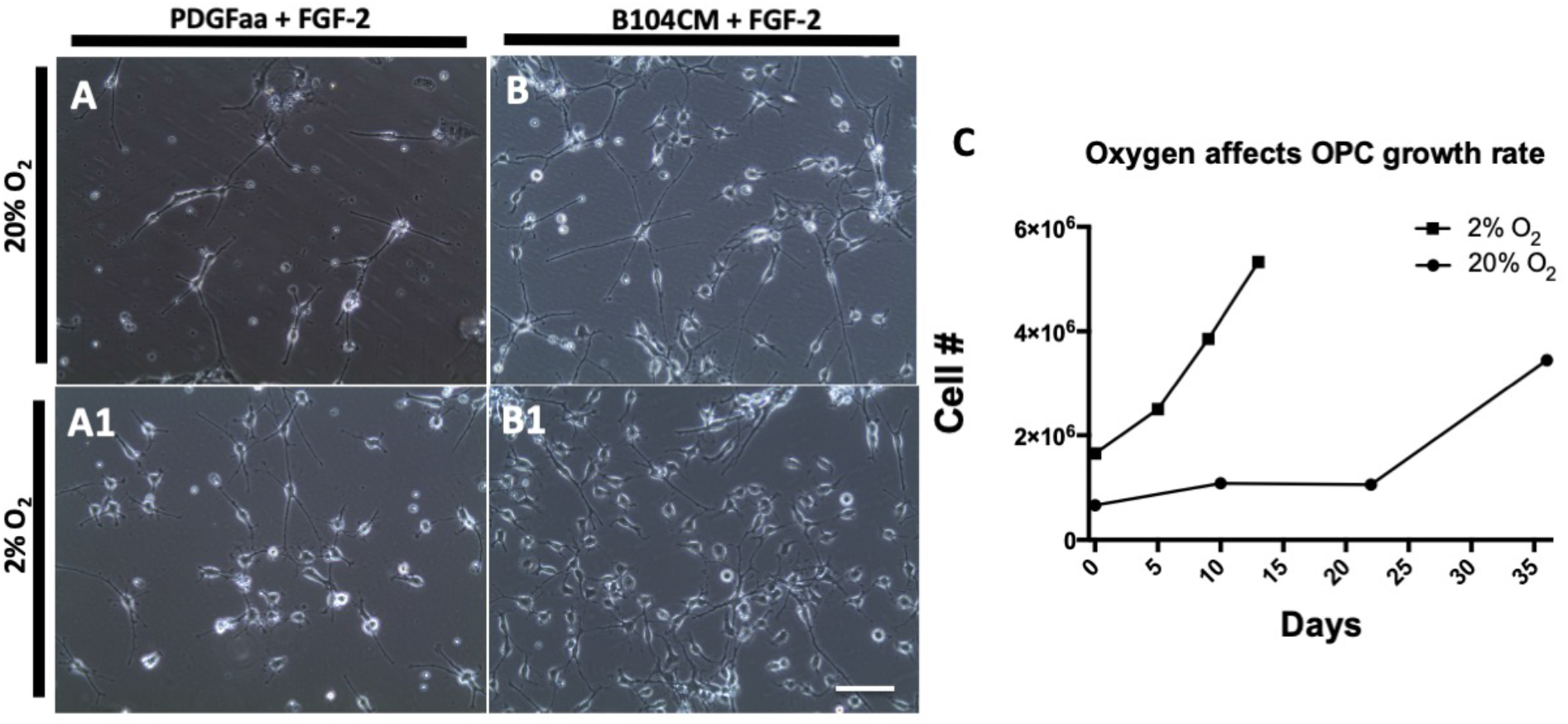
Murine OPC growth is accelerated when cultured in medium supplemented with B104CM and FGF2 and grown in 2% O_2_. The enriched OPC fraction was plated onto fibronectin coated tissue culture plates at 3×10^4^ cells/cm^2^ immediately after the second shake and cultured in B104CM+FGF2 or PDGF-aa+FGF-2 in 20% Oxygen (A, B) or 2% Oxygen (A1, B1) for 5 days. The number of cells obtained when grown in B104CM+FGF2 in either 2% or 20% Oxygen over 4 serial passages across 35 days is depicted (C). Scale bar represents 50μm.

It has been reported that both human and mouse neural progenitors grow better when maintained in physiologically oxygenated medium [38,39]. Therefore, the OPCs were cultured in a 3-gas incubator with humidified 2% oxygen, 5% carbon dioxide and 97% nitrogen. Cells were cultured in medium containing either 10 ng/mL PDGFaa + 10 ng/mL FGF-2 or 30% B104CM + 10 ng/mL FGF-2 (Figure 2A1, B1). OPCs grew at a much faster rate when cultured in B104CM + FGF-2 at 2% oxygen than when cultured in FGF-2 and PDGF, and they could be passaged more frequently than cells maintained in the same medium but propagated in 20% oxygen (Figure 2C). Cells maintained in 20% oxygen, if let alone for an extended period of time, did expand but their numbers never reached the cell numbers obtained in a 2% oxygen incubator. Growth in 2% oxygen significantly increased the overall yield of cells obtained such that by passage 3, 4×10^6^ cells were obtained, whereas it took another passage and over 10 days for this number of cells to be obtained when the cells were maintained in 20% oxygen.

### OPCs are overwhelmingly A2B5+ and Ki67+ when grown in B104+FGF2 in physiological oxygen

Next we assessed the effects of cell density on OPC proliferation. OPCs were passaged twice and then plated onto fibronectin coated chamber slides at 3×10^4^ cells/cm^2^(approximately 25% confluency) (Figure 3A,B) and at 6×10^4^ cells/cm^2^(approximately 50% confluency) (Figure 3C,D). The cells were maintained for 4 days in a 2% oxygen incubator and then stained for markers of the oligodendrocytic lineage (A2B5 and Olig2). OPCs maintained their early progenitor phenotype with the majority of cells expressing A2B5 (Fig. 3A,C). When the percentage of cells double positive for the cell proliferation marker Ki67 and Olig2 (Fig. 3B,D) was quantified to produce a mitotic index, the mitotic index was not significantly different between the two plating densities conditions and was equivalent to the number of Olig2+ cells (Figure 3E).

**Figure 3.**
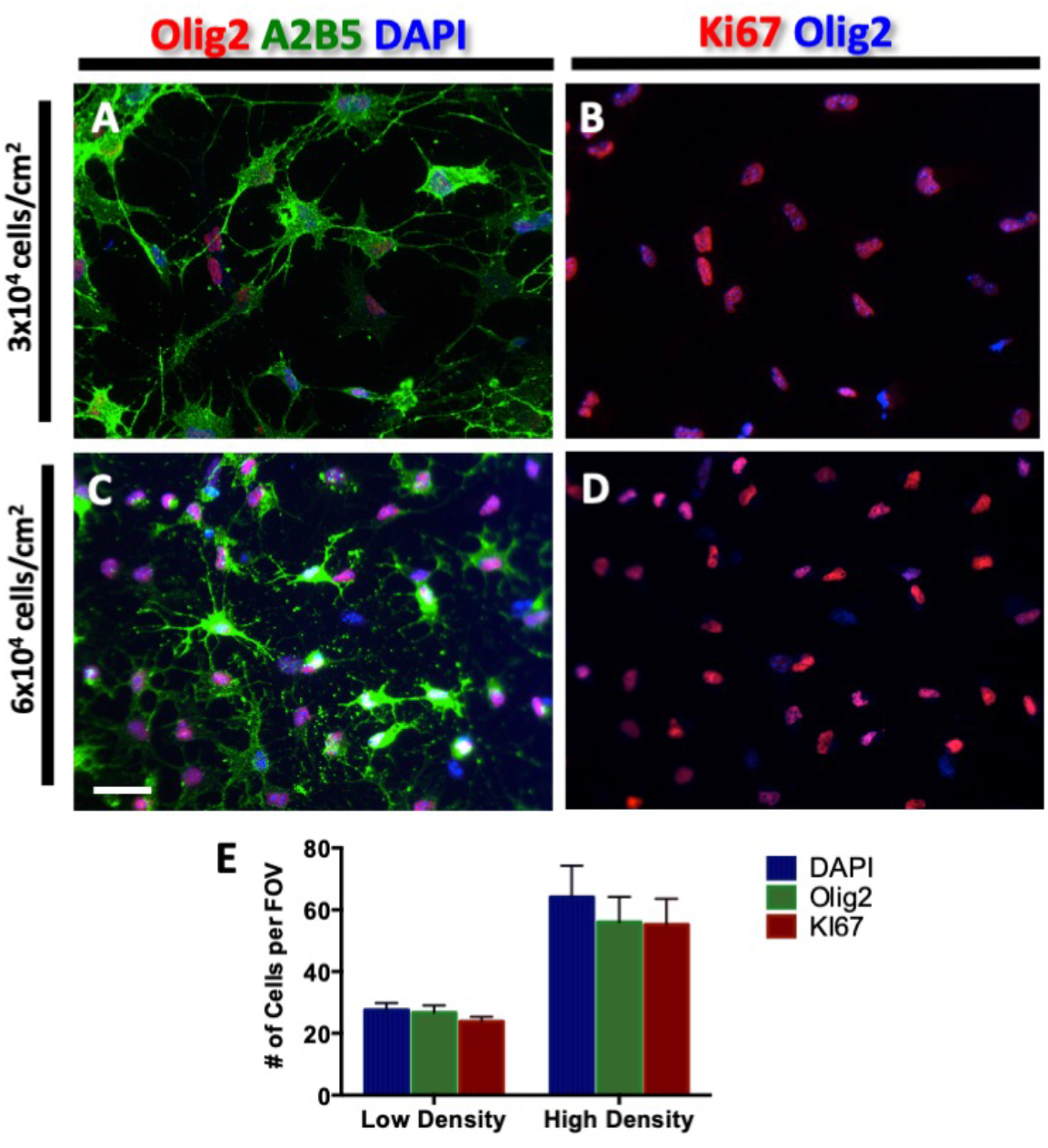
OPCs are A2B5+ and Ki67+ at both low and high density when grown in B104CM+FGF2 in 2% oxygen. After the first passage, OPCs were plated onto fibronectin coated chamber slides at 3×10^4^ cells/cm^2^ (A) or 6×10^4^cells/cm^2^ (B) and maintained for 4 days in medium supplemented with B104CM+FGF2 in a 3 gas incubator with 2% O_2_, 5% C0_2_ and 93% N_2._ (**A**,**B)** Cells were stained for A2B5 (green), Olig2 (red) and DAPI (blue). (A1Olig2-blue and Ki67-red (A1, B1). The number of DAPI+, Olig2+ and Ki67+ cells was quantified (C). Scale bar represents 50μm. Data represent mean ± SEM from 3 independent experiments.

### Mitogens have differential effects on the phenotype of cultured OPCs

As OPCs mature, they lose A2B5 expression and acquire markers associated with later stage progenitors, such as O4. To determine whether the combination of PDGF and FGF-2 would prevent mouse OPCs from maturing they were cultured in PDGFaa+FGF-2 or FGF-2 for 94 h after the second shake. At 24 h, in both conditions, the number of NG2+ cells was equivalent, but after 96 h the number of cells in PDGFaa+FGF-2 had increased significantly whereas the number of cells cultured with FGF-2 alone had not significantly increased from those initially plated (Figure 4A). Similar to rat OPCs cultured in PDGFaa+FGF-2, most of the cells were A2B5+ (90%±0.03), 8%±0.01 expressed O4 and there was only a small fraction (2%) that were GFAP+ (Figure 4B, D, D1 D2). Of those cells cultured in FGF-2 alone, a greater percentage started to express O4 (19%±0.04) (Figure 4B, 4C, C1, C2). However, a large fraction of the cells maintained in FGF-2 alone were negative for both A2B5 and O4 (∼40%). These cells stained for Olig2 (Figure 4C1) and unexpectedly GFAP (Figure 4C2).

**Figure 4.**
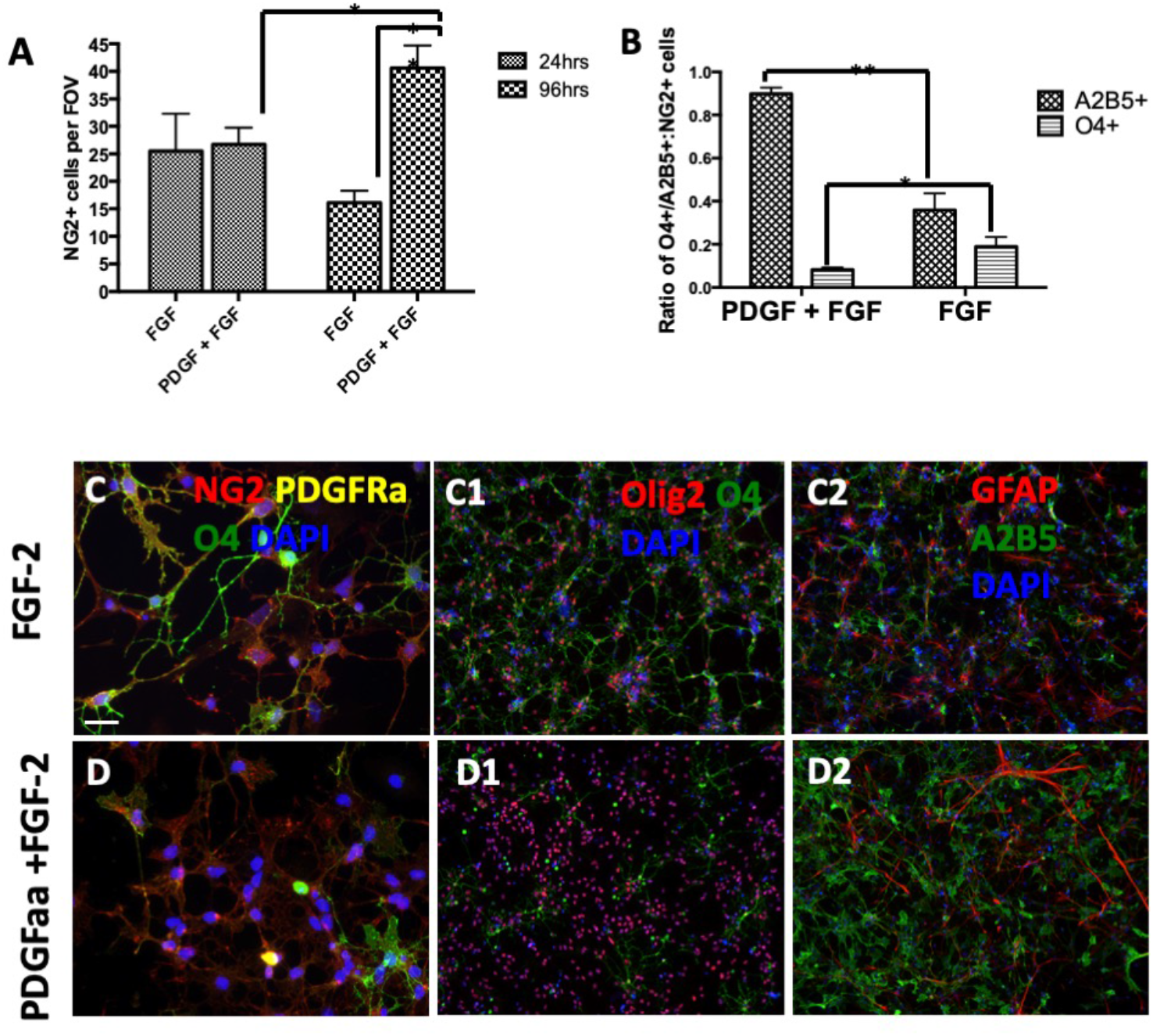
Differential effects of mitogen combinations on the phenotype of cultured OPCs. OPCs were plated onto fibronectin coated chamber slides at 3×10^4^ cells/cm^2^ in media supplemented with FGF2 or PDGF-aa+FGF-2 and maintained in 2% O_2_. The number of NG2+ cells was counted at 24 and 96 h after initial plating (A). The ratio of NG2+ cells that were positive for A2B5 or O4 was quantified at 96 h after plating (B). Data represent mean ± SEM from 3 independent experiments. * = *p* < 0.05, ** = *p* < 0.01 by student’s *t* test. Cells cultured in FGF-2 or PDGF-aa+FGF-2 were stained with NG2 (red), PDGFRα (yellow), O4 (green) and DAPI (blue) (C, D). Cells were stained for for O4 (green) Olig2 (red), DAPI (blue) (C1, D1) or GFAP (red), A2B5 (green) and DAPI (blue) after 4 days in culture (C2, D2). Scale bar represents 20μm (C, D) and 200um (C1, C2, D1, D2).

### OPCs progress through the oligodendrocytic lineage overtime when grown in B104+FGF-2 and 2% O_2_

Whereas most immunofluorescence studies are limited to analyzing 4 fluorescent channels, flow cytometry can simultaneously analyze upwards of 13 markers and we have found that at least 5 fluorophores are required to distinguish stem cells from multipotential progenitors [40,41]. Therefore, to better characterize the cell types present within these OPC enriched cultures we used flow cytometry to analyze the expression of 7 surface markers (Supplementary Table 1). The markers chosen assessed several stages of OPC maturation as well as markers associated with neural stem cells, bipotent, and multipotent progenitors. Ninety nine percent of the cells propagated in B104CM and FGF2 for 3-6 passages expressed NG2 and PDGFRα (Figure 5A, B). None expressed the phenotype of a neural stem cell marker CD133+/LeX+ (Figure 5A1, B1). However, whereas only 17% of the cells expressed O4 at passage 3, expression of O4 increased to 98% by passage 6 (Figure 5A2, B2). Additionally, initially only a few of the OPCs expressed Glast when cultured in B104CM+FGF2 (Figure 5A2, B2); however, by passage 6, 68% of the OPCs expressed Glast, and most of these cells were also O4+.

**Figure 5.**
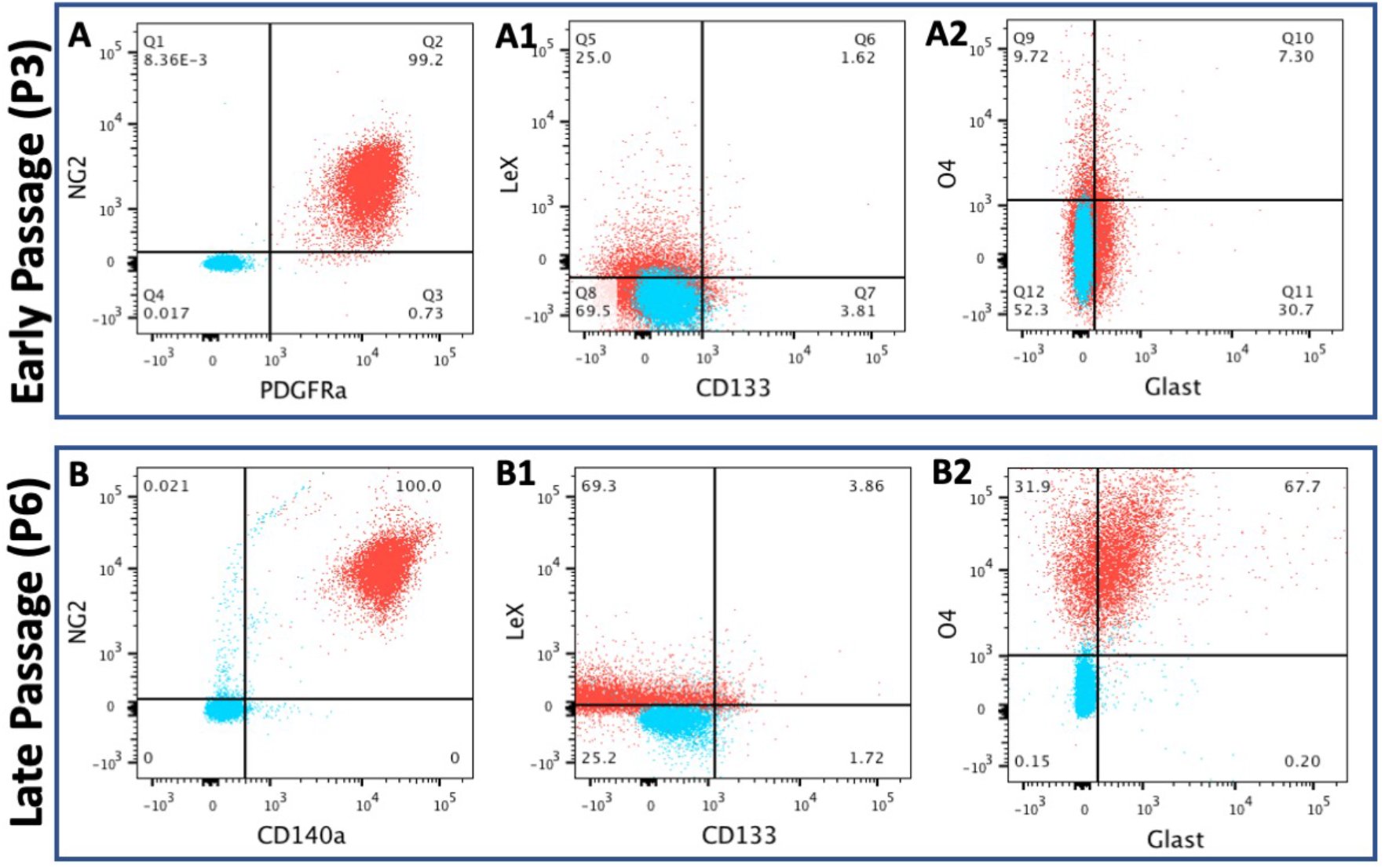
OPCs acquire late progenitor cell markers with time in culture when grown in B104CM+FGF2 and 2% O_*2*._ OPCs were grown *in vitro*, passaged, stained and then analyzed live by Flow Cytometry at passage 3 (A-A2) and passage 6 (B-B2) using a flow panel comprised of CD133/LeX/NG2/CD140a/O4/Glast. 50, 000 events were analyzed per group. Forward and side scatter were used to exclude debris and DAPI exclusion was used to define live cells. Blue dots represent isotype controls for background and gating standards while red dots represent the stained events. Data are representative of two independent experiments.

### Some OPCs can differentiate into mature myelinating oligodendrocytes in vitro when treated with T3, CNTF and TGFβ1

The presence of B104CM and FGF2 has been used to maintain OPCs in a primitive proliferating state and removing these mitogens, should promote their terminal differentiation into myelinating oligodendrocytes. To determine whether these mouse OPCs could mature OPCs from passages 1 and 3 were detached and plated onto laminin coated dishes. They were maintained in FGF-2 for 24 h to allow them to gradually mature at which time all mitogens were removed. They were then maintained in a basal medium supplemented with XXXX LISA, PLEASE DESCRIBE FIGURE 6.

**Figure 6.**
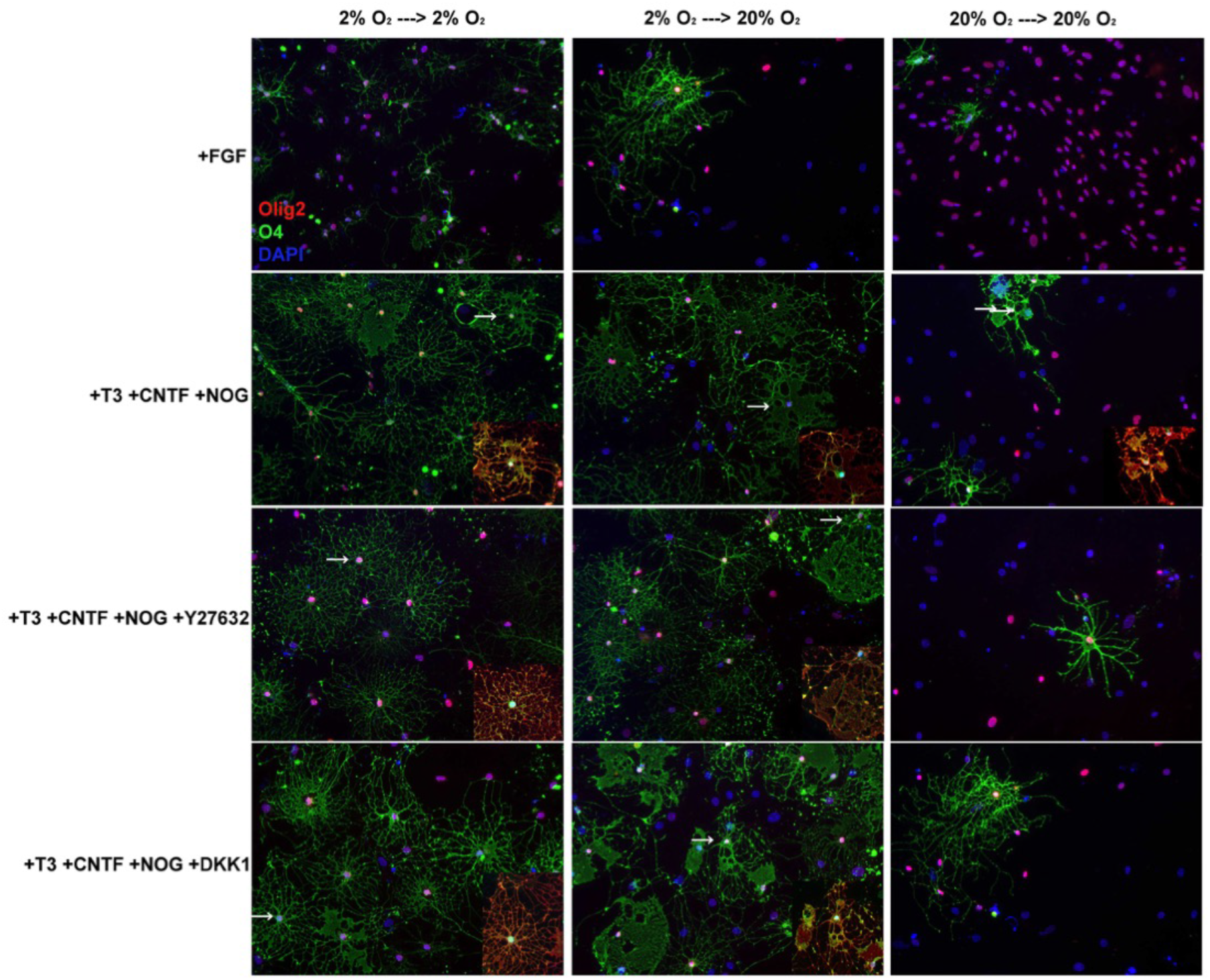
Effects of culture supplements and oxygen on differentiation of early passage OPCs. OPCs after 1 passage that had been propagated in B104CM+FGF2 and were plated onto poly-d-lysine and laminin coated chamber slides in FGF2 for 24 hours. Mitogens were removed and media was replaced with N2B2 supplemented with either FGF-2, T3+CNTF+noggin, T3+CNTF+noggin +Y27632, or T3+CNTF+noggin+DKK1. Cells were maintained for 7 days and then stained for O4 (green), Olig2 (red) and DAPI (blue). Scale bar represents 50μm

In parallel, OPCs from passage 6 were differentiated in combinations of 1.95 μg/mL T3, 1 ng/mL rrCNTF and 1 ng/mL TGFβ1. The cells were maintained in a 2% O_2_ incubator with medium changes every 2-3 days for 7 days. Under these conditions, cells expressing O4, PLP, MBP, or O1 were rarely seen with T3 supplementation alone (Figure 7A, A1). By contrast when the medium was supplemented with either T3+CNTF or T3+TGFβ1, many more cells could be found that expressed the myelin proteins MBP and PLP and these cells were surrounded by multiple, sheet-like membranes (Figure 7B, B1, C, C1). However, the CNTF addition increased the relative percentage of cells expressing GFAP+ (Figure 7B1).

**Figure 7.**
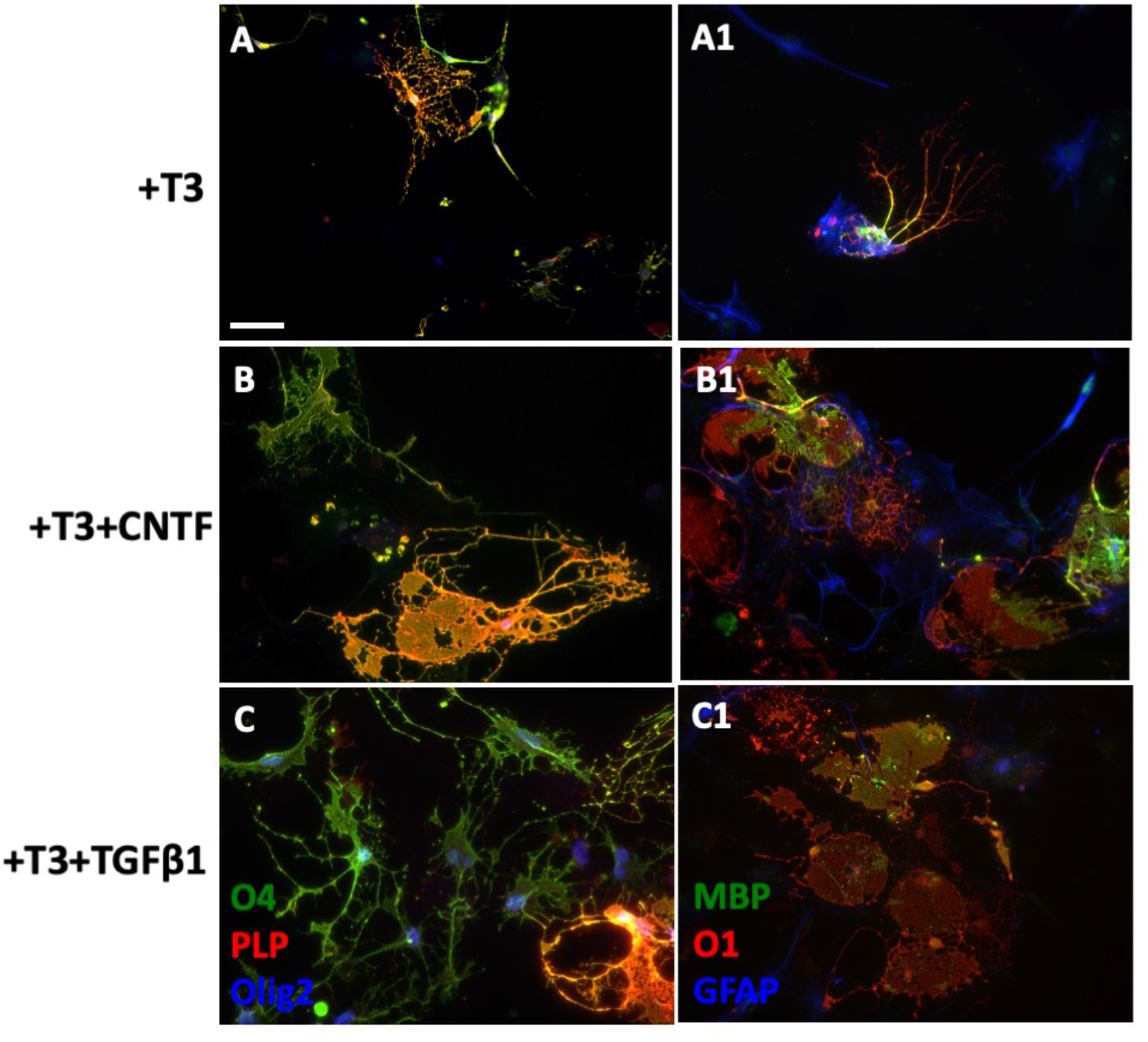
Effects of culture supplements and oxygen on differentiation of late passage OPCs. OPCs were grown in B104CM+FGF2 for 6 passages and then plated onto poly-d-lysine and laminin coated chamber slides in FGF2 for 24 hours. Mitogens were removed and media was replaced with N2B2 supplemented with either T3, T3+1 ng/mL CNTF or T3+1 ng/mL TGFβ1. Cells were maintained for 7 days and then stained for O4 (green), PLP (red), Olig2 (blue) (A, B, C) or MBP (green), O1 (red), GFAP (blue) (A1, B1, C1). Scale bar represents 50μm

## Discussion

Our current knowledge of OPCs has largely been derived from studies on rat OPCs. Rat OPCs from embryonic, neonatal and adult CNS can be isolated and expanded *in vitro* for more than a year in the presence of platelet derived growth factor aa (PDGF-aa) and fibroblast growth factor 2 (FGF-2) [42,43]. In contrast to the rat OPC cultures, relatively few studies have been reported using highly enriched mouse OPC cultures, despite the fact that the mouse is the animal of choice for rodent models of demyelinating diseases [22,29,44-47]. Quite simply it has been technically challenging to obtain highly homogeneous cultures of mouse OPCs *in vitro* in large numbers. This study establishes a simple method to purify and expand large numbers of mouse OPCs from the brains of neonatal mice *in vitro*.

Initially, we followed the well-established protocol described by McCarthy and de Vellis [23] to separate the OPCs from microglia, astrocytes and fibroblasts. During these early experiments, we discovered differences in numbers of OPCs produced in the mixed glial cultures that correlated with different lots of FBS. Therefore, to maximize OPC production during the early stages of the *in vitro* culture, we screened several lots of FBS from different commercial suppliers. The serum chosen for future use was based on the numbers of OPCs growing on top of the bedlayer. A dozen bottles of this lot was purchased and used for all subsequent experiments.

There were notable differences in survival and proliferation rate of mouse vs rat OPCs. Since Bottenstein and her colleagues (1988) discovered preferential proliferation of rat OPCs in B104 neuroblastoma conditioned medium (B104CM), B104CM has been widely used to generate highly enriched primary rat OPC cultures [48,49]. Subsequent studies have confirmed the presence of PDGF-aa and neuregulin 1 in B104CM — mitogens that influence OPC growth [50,51]. Therefore, we compared the growth of the OPCs in medium supplemented with B104CM to medium supplemented with PDGF-aa. We observed a significant increase in the expansion of OPCs grown in B104CM+FGF2. It was previously reported that a greater number of O4+ late stage OPCs were observed in mouse OPC cultures following isolation from the brain tissue than seen in the rat OPC cultures [17]. This may indicate that initially the mouse OPCs that are isolated are at a more advanced developmental stage and are less proliferative. Therefore, PDGF-aa and FGF2 alone are not sufficient to drive their expansion whereas the B104CM contains some unidentified factor that causes these cells to dedifferentiate to an early stage OPC and to proliferate at a greater rate. This factor may act in a species-specific way since this phenomenon is not noted in rat OPC cultures.

Another major difference in our method for culturing mouse OPCs was to decrease their exposure to oxygen. Our previous studies have shown that neural progenitor cell proliferation and differentiation can be affected by growth in 20% vs. 2% oxygen incubators [52]. Yet other studies have shown that oxygen tension is a critical determinant in NSC propagation whereas growth in 5% O_2_ expanded Nestin+/CD133+/CD24+ precursors that readily differentiated into all three neural lineages [38,39]. Therefore, we evaluated the effect of oxygen concentration on mouse OPCs over 4 passages. These studies revealed a major difference in cell proliferation and survival when OPCs were expanded in 2% oxygen. Large cells numbers could be obtained rapidly and the cells maintained their expression of A2B5/Olig2/Ki67 over several passages. However, additional studies will need to be conducted to determine whether the differences are attributable to proliferation or cellular death. Additionally, it remains possible that the culture conditions are selecting for a specific clone or there is a population shift as opposed to a true maturation event.

An earlier study had concluded that cAMP addition was required for mouse OPCs to survive in B104CM+FGF2 [17]. However, cAMP addition was not required in our studies. A plausible hypothesis is that the cAMP or downstream targets of protein kinase A counter-act the toxic effects of oxidative stress induced by growth in superphysiological oxygen – however, we did not test this hypothesis. To understand the molecular changes responsible for the difference in growth, RNA was collected from OPCs cultured in 20% oxygen and 2% oxygen. These samples are being assessed by RNASeq technology and the results are pending.

Many labs have struggled with mouse OPCs differentiation *in vitro* [18,22], and we faced similar challenges. When the mitogens were removed and cells were placed into differentiation medium that was supplemented with T3 there was a reduction in the proportion of cells of the oligodendrocytic lineage and an increase in cells of the astrocytic lineage. It’s possible that the presence of contaminating astrocytes will expand under certain conditions. When we removed PDGF-aa from the media and only supplement it with FGF2 we noticed a higher number of O4+ OPCs but we also detected an increased number of GFAP+ astrocytes. Since astrocytes express FGFR-2 [53] the presence of FGF2 likely caused an expansion of contaminating astrocytes.

In order to assess the mouse OPCs capacity to fully differentiate into a terminal myelinating oligodendrocyte, we tested the effects of additional factors besides T3 that had been tested on rat OPC cultures. Unlike rat OPCs that easily differentiate into mature oligodendrocytes when all of the mitogens are removed and the media is supplemented with T3 [54], mouse OPCs do not behave in the same manner. T3 alone was not sufficient for OPC differentiation. In fact, most of the oligodendrocyte lineage cells did not survive in this condition. OPC maturation was enhanced by the addition of either CNTF or TGFβ1. Medium supplemented with CNTF increased the proportion of PLP+ O1+ and MBP+ oligodendrocytes, and these cells had highly complex arborizations. However, a considerable number of differentiated astrocytes were observed which supports a previously described role for CNTF [55-57]. This suggests that CNTF may be playing a synergistic role with soluble factors in the media to enhance OPC differentiation, but further investigation is needed to determine whether greater differentiation along the oligodendroglial lineage in the presence of CNTF is possible. The addition of TGFβ1 also increased OPC maturation into PLP+, O1+ oligodendrocytes with MBP immunoreactivity and minimal GFAP+ astrocyte formation consistent with earlier studies on rat OPCs that found that TGFß1 reduced OPC cell proliferation and increased their maturation [30]. Since very few OPCs fully differentiated into myelin producing oligodendrocytes, the media likely lacked all the tropic factors necessary for maturation. It has been shown that BDNF synthesized and released from astrocytes during a demyelinating lesion enhances proliferation and differentiation of endogenous NG2+ cells [58-60]. Therefore, in future studies we will test the effects of BDNF on our cultured OPCs.

Multiple approaches have been developed to purify OPCs, including immunopanning and oligosphere formation. Each method has its own benefits and complications. For example, mouse OPCs produced from oligospheres involves neurosphere formation from embryonic mouse brains followed by oligosphere induction [22,61]. While this method produces oligodendrocytes, it is not clear that they are identical to those formed from postnatal brains since they are derived from stem cells as opposed to primary OPCs. Immunopanning of postnatal mouse brains has also been utilized; however, these OPCs are usually selected by A2B5 monoclonal antibody, and this antigen may not select exclusively for cells that will differentiate within the oligodendroglial lineage [62]. A2B5+ OPCs have also been enriched from postnatal mouse brains by magnetic cell sorting [18], but the yield is very low. Moreover, it has been suggested that these OPCs are limited to 2 passages before the OPCs will differentiate into type 2 astrocytes [18], which is highly impractical for studying these cells. In our studies, we were able to passage these cells for up to 10 times.

While we were able to passage the mouse OPCs extensively, they did change with time in culture. More specifically, they acquired the O4 antigen as described in rat cultures [63], as well as increased expression of GLAST. This result is reminiscent of those performed by Mark Noble who found that with repeated passage of early O-2A cells that they became more like adult O-2A cells [2]. Adult O-2As differ from early O-2As in several respects for example their cell cycle, whereas perinatal cells have a short 18-hour cell cycle as compared to an adult’s 65 hour cycle length. Perinatal cells migrate rapidly (21 μm/h) while adult cells migrate slowly (4.3 μm/h). Perinatal cells differentiated rapidly, synchronously and symmetrically while adult cells differentiate slowly and asymmetrically *in vitro* [2,32,42,64-66]. Thus, for investigators interested in studying the adult OPCs, our method of repeatedly passaging the OPCs will enable them to produce large numbers of cells for analysis relatively simply, whereas acquiring similar numbers of cells directly from the adult brain would be prohibitively complex.

Overall, compared to other mouse OPC isolation methods, the method that we have described here is simple and straightforward. Our results demonstrate that there is a quantitative difference between mouse OPCs grown in PDGF-aa+FGF-2 vs. B104CM+FGF-2 and also that mouse OPCs are very sensitive to 2% O_2_ vs. 20% O_2_. This method can serve as an improved *in vitro* model for the study of primary mouse OPCs in development, disease and trauma.

## Conclusion

A method that is reproducible, simplistic and can easily expand large numbers of mouse OPCs is desirable for many researchers who perform experiments on primary cells and transgenic mice models. In order to optimize this procedure, we determined the necessary *in vitro* environment for rapid and sustainable mouse OPC expansion. In these studies, we have shown mouse OPCs can be easily fractionated from other glial cells and expanded in culture for an indefinite period of time. The growth of mouse OPCs, unlike rat OPCs, is inhibited by superphysiological levels of oxygen and therefore oxygen concentration can have a major effect on cell expansion. And lastly with time in culture mouse OPCs acquire the phenotype of adult O-2A progenitors and thus behaves as the cells would normally *in vivo*.

## Figures and tables

**Supplementary Figure 1.**
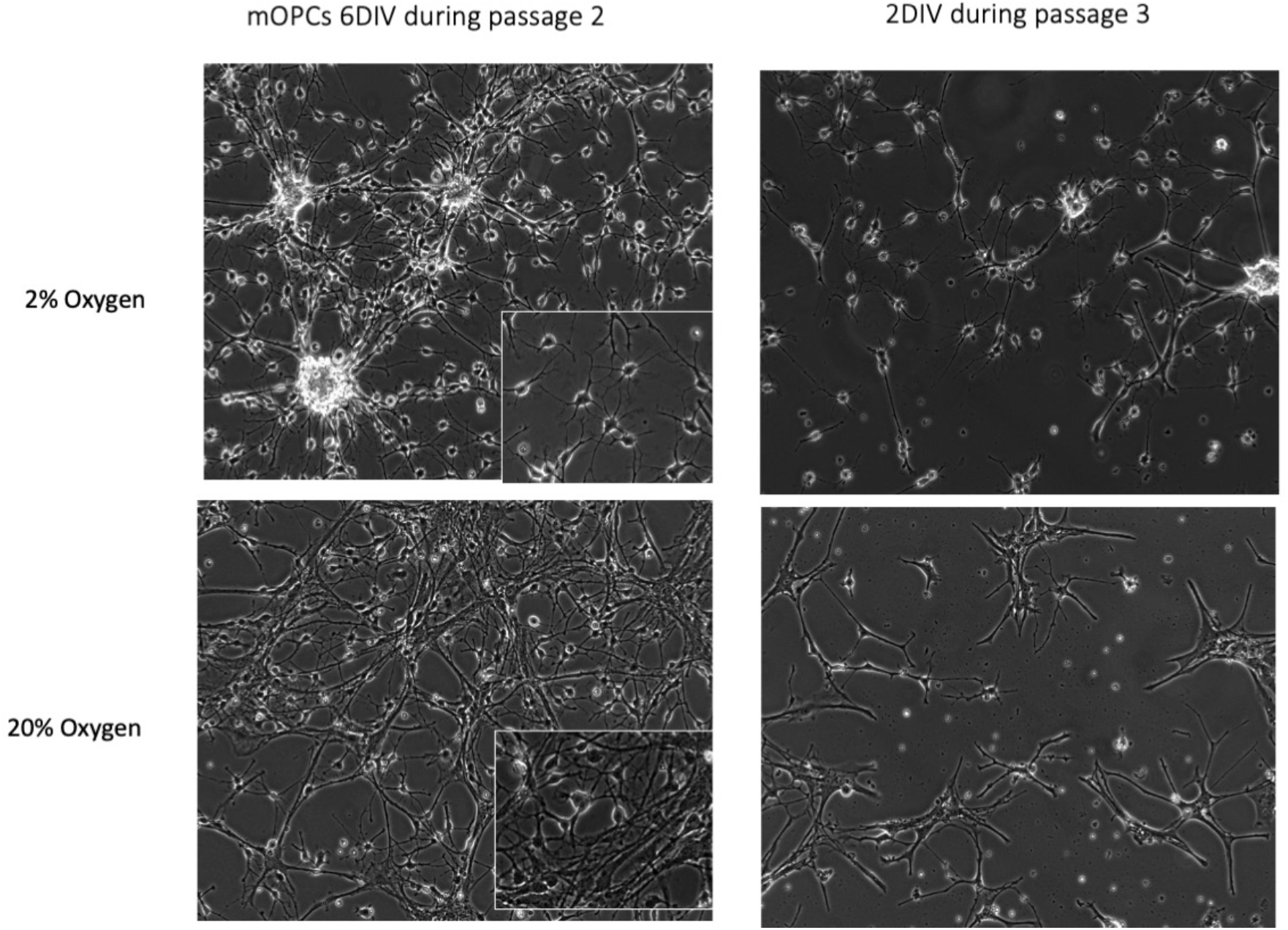

**Supplementary Table 1.**

| **Antibody** | **Antigen and/or AKA** | **Targets** |
| --- | --- | --- |
| CD133 | Prominin-1 | Stem cells and Ependymal cells |
| CD15 | Lewis-X, LeX, SSEA-1 | Neural precursors, some immature astrocytes |
| NG2 | Chondroitin sulfate<br>proteoglycan | Multipotential, bipotential and unipotential progenitors |
| CD140a | PDGFRα | Multipotential, bipotential and unipotential progenitors |
| O4 | Sulfatide, POA | Late OPCs |
| A2B5 | Complex Gangliosides | Early OPCs and immature neural precursors |
| Glast | Glutamate Transporter,<br>EEAT1 | NSCs, astrocytes and cerebellar oligodendrocytes |

